# Tract-specific white matter microstructure alterations among young adult *APOE* ε4 carriers: A replication and extension study

**DOI:** 10.1101/2022.03.23.485532

**Authors:** Rikki Lissaman, Thomas M. Lancaster, Greg D. Parker, Kim S. Graham, Andrew D. Lawrence, Carl J. Hodgetts

**Author notes:** Corresponding author: Carl J. Hodgetts. These authors jointly supervised this work (Andrew D. Lawrence, Carl J. Hodgetts).

## Abstract

The parahippocampal cingulum bundle (PHCB) connects regions known to be vulnerable to early Alzheimer’s disease (AD) pathology, such as posteromedial cortex and medial temporal lobe. While AD-related pathology has been robustly associated with alterations in PHCB microstructure, specifically lower fractional anisotropy (FA) and higher mean diffusivity (MD), emerging evidence indicates that the reverse pattern is evident in younger adults at-risk of AD. In one such study, Hodgetts et al. (2019) reported that healthy young adult carriers of the apolipoprotein-E (*APOE*) ε4 allele – the strongest common genetic risk factor for AD – showed higher FA and lower MD in the PHCB but not the inferior longitudinal fasciculus (ILF). These results are consistent with proposals claiming that heightened neural activity and connectivity have a significant role in posteromedial cortex vulnerability to amyloid-β and tau spread beyond the medial temporal lobe. Given the implications for understanding AD risk, here we sought to replicate Hodgetts et al.’s finding in a larger sample (*N* = 128; 40 *APOE* ε4 carriers, 88 *APOE* ε4 non-carriers) of young adults (age range: 19-33). Extending this work further, we also conducted exploratory analyses using a more advanced measure of microstructure: hindrance modulated orientational anisotropy (HMOA). These analyses included an investigation of hemispheric asymmetry in PHCB and ILF HMOA. Contrary to the original study, we observed no difference in PHCB microstructure between *APOE* ε4 carriers and non-carriers. Bayes factors (BFs) further revealed moderate-to-strong evidence in support of these null findings. *APOE* ε4-related differences in ILF HMOA asymmetry were evident, however, with carriers demonstrating lower leftward asymmetry. Our findings indicate that young adult *APOE* ε4 carriers do not show alterations in PHCB microstructure, as observed by Hodgetts et al., but may show altered asymmetry in ILF microstructure.

## 1. Introduction

Alzheimer’s disease (AD) is a chronic, progressive disease and the most common cause of dementia (Scheltens et al., 2021). The hallmark pathological features of AD are the presence of extracellular amyloid-β-containing plaques and intracellular tau-containing neurofibrillary tangles (DeTure & Dickson, 2019; Trejo-Lopez et al., 2021). Although controversial (Frisoni et al., 2022; Herrup, 2015), the dominant hypothesis in the field – the amyloid cascade hypothesis – holds that the accumulation of amyloid-β peptide is the critical factor in AD pathogenesis (Selkoe & Hardy, 2016). Amyloid-β accumulation follows a relatively distinct spatiotemporal pattern in the ageing brain, beginning preferentially in posteromedial regions, including retrosplenial/posterior cingulate cortices and precuneus (Mattsson et al., 2019; Palmqvist et al., 2017; Villeneuve et al., 2015). Collectively, these regions are sometimes referred to as posteromedial cortex (Parvizi et al., 2006). The vulnerability of posteromedial cortex to AD pathology has been linked to its hub-like properties (Jagust, 2018), in particular its high-levels of baseline metabolic/neural activity and high intrinsic/extrinsic connectivity (Bero et al., 2012; Buckner et al., 2009; de Haan et al., 2012). Notably, posteromedial cortex is densely connected with several medial temporal lobe structures, such as parahippocampal cortex and hippocampus, forming a “posterior medial” or “extended navigation” network (Murray et al., 2017; Ranganath & Ritchey, 2012). This broader network is implicated in several inter-related cognitive functions that are impaired early in AD, such as episodic memory (Rajah et al., 2017), perceptual scene discrimination (Lee et al., 2006), and spatial navigation (Coughlan et al., 2018). Given this, there is a pressing need to identify biomarkers that capture the functional and/or structural integrity of this AD-vulnerable brain network. In this context, the parahippocampal cingulum bundle (PHCB) – a prominent white matter tract linking posteromedial cortex with the medial temporal lobe (Bubb et al., 2018; Heilbronner & Haber, 2014; Jitsuishi & Yamaguchi, 2021) – represents a strong candidate for understanding and characterising connectivity alterations associated with AD.

Increasing evidence indicates that PHCB connectivity is altered in AD. Using diffusion magnetic resonance imaging (dMRI), a non-invasive method that examines the random, microscopic movement of water molecules, it is possible to delineate the major white matter tracts of the brain and evaluate their microstructural properties in vivo (Assaf et al., 2019). In most AD-relevant dMRI studies, white matter microstructure is assessed via measures derived from the diffusion tensor, notably fractional anisotropy (FA) and mean diffusivity (MD; Harrison et al., 2020). Low FA and high MD are widely interpreted as representing poorer microstructural integrity and thus lower connectivity (Yeh et al., 2021), although multiple biological factors – including neuroinflammation (Kor et al., 2022) – can influence these measures (Jones, Knösche, & Turner, 2013). Studies comparing AD patients to cognitively normal older adults using dMRI have reliably observed both lower FA and higher MD in the cingulum bundle and the PHCB in particular (Acosta-Cabronero et al., 2010; Bozzali et al., 2012; Choo et al., 2010; Kantarci et al., 2017). In addition, longitudinal changes in PHCB microstructure – reduced FA, increased MD – have been reported among AD patients but not cognitively normal older adults (Mayo et al., 2017). Indeed, it has recently been suggested that PHCB FA constitutes a highly effective biomarker for differentiating between typical ageing and AD (Dalboni da Rocha et al., 2020).

Studies of amnestic mild cognitive impairment (aMCI), a transitional stage between typical ageing and AD (Albert et al., 2011), further highlight that PHCB alterations precede the onset of AD dementia. In one region-of-interest (ROI) meta-analysis, for example, Yu et al. (2017) identified robust alterations in PHCB microstructure (lower FA, higher MD) among individuals with aMCI. This is congruent with the notion that cingulum bundle alterations predict cognitive decline in aMCI and may even predict conversion to AD (Gozdas et al., 2020). Studies combining positron emission tomography and dMRI have also allowed PHCB changes to be linked directly to AD pathology. For example, amyloid-β burden has been associated with longitudinal changes in white matter microstructure that are consistent with patterns observed in aMCI and AD (Rieckmann et al., 2016; Song et al., 2018; Vipin et al., 2019). In particular, high levels of cortical amyloid-β burden at baseline have been associated with accelerated decline in PHCB FA and a trend-level increase in PHCB MD (Rieckmann et al., 2016). In keeping with this tract-specific finding, one recent cross-sectional study reported that lower FA and higher MD in the PHCB was associated with greater cortical amyloid-β and entorhinal tau burden, especially in those with high levels of pre-existing pathology (Pichet Binnette et al., 2021). It thus appears that PHCB microstructure is detrimentally impacted over the course of AD, including stages prior to the onset of dementia symptoms.

Emerging research indicates, however, that asymptomatic individuals exhibit alterations in white matter microstructure that run counter to the characteristic AD pattern. Illustrating this point, several studies have observed higher FA and lower MD in early-stage amyloid-β pathology, a pattern that is reversed as pathology further accrues (Collij et al., 2021; Dong et al., 2020; Wolf et al., 2015). These findings point to a biphasic pattern of microstructure over the disease course, with a period of high FA/low MD preceding the pattern commonly observed in patients with aMCI and AD. While increased FA in the context of early AD pathology could reflect neuroinflammation (Benitez et al., 2021; Dong et al., 2020), there is evidence that heightened activity and connectivity – including structural connectivity – may actually precede AD pathology, predisposing individuals to later amyloid-β deposition (Bero et al., 2012; Buckner et al., 2009; de Haan et al., 2012). Support for this proposal can be found in studies of young adults carriers of the apolipoprotein-E (*APOE*) ε4 allele. The *APOE* ε4 allele is the strongest common genetic risk factor for AD (Belloy et al., 2019), and is also associated with a younger age of onset and faster rate of posteromedial amyloid-β accumulation (Burnham et al., 2020; Mishra et al., 2018). In line with the notion that this amyloid-β accumulation is related to earlier connectivity changes, a study applying graph theoretical analysis to dMRI data observed that age was negatively associated with local interconnectivity in posteromedial regions, but only among *APOE* ε4 carriers (Brown et al., 2011). Higher levels of local interconnectivity in younger adults drove this finding, such that there was a putative *APOE* ε4-related increase in connectivity early in life that was subsequently followed by a sharper decline later in the lifespan (Brown et al., 2011; see also Ma et al., 2017). Relatedly, Felsky and Voineskos (2013) further reported higher cingulum bundle FA in younger *APOE* ε4 carriers compared to younger non-carriers, but lower cingulum bundle FA in older *APOE* ε4 carriers compared to older non-carriers. Given that young adults are unlikely to possess significant amyloid-β burden (Jansen et al., 2015), these findings suggest that early-life structural alterations may precede pathology.

Consistent with this, Hodgetts et al. (2019) observed higher FA and lower MD among *APOE* ε4 carriers relative to non-carriers in the PHCB but not the inferior longitudinal fasciculus (ILF), a tract that connects the occipital lobe to the ventro-anterior temporal lobe (Herbet et al., 2018). Hodgetts et al. also found that PHCB microstructure was correlated with posteromedial cortex activity during perceptual scene discrimination, a task that has previously been shown to elicit heightened activity in young *APOE* ε4 carriers (Shine et al., 2015) and is sensitive to AD (Lee et al., 2006). Based on the proposal that heightened neural activity and connectivity can have a significant role in hub-like vulnerability to amyloid-β (Bero et al., 2012; Buckner et al., 2009; de Haan et al., 2012), it is plausible that such early-life PHCB alterations may explain why *APOE* ε4 is associated with earlier and faster posteromedial amyloid-β accumulation (Burnham et al., 2020; Mishra et al., 2018). Moreover, as the spread of tau has been linked to heightened functional connectivity between posteromedial cortex and the medial temporal lobe (Ziontz et al., 2021) – presumably mediated by the PHCB (Jacobs et al., 2018) – it is possible that early-life increases in structural connectivity are also related to elevated tau in *APOE* ε4 carriers (Therriault et al., 2020).

In view of the potential implications for understanding the role of *APOE* ε4 in AD risk, we sought to replicate Hodgetts et al.’s (2019) finding that healthy young adult *APOE* ε4 carriers demonstrate higher FA and lower MD than non-carriers in the PHCB but not the ILF. We analysed data from an independent data set of young adults, with a total sample over four times larger than the original study. This replication attempt thus constitutes an important test of the notion that increased PHCB connectivity, as indexed by higher FA and lower MD, is evident in young adult *APOE* ε4 carriers, potentially increasing vulnerability to both amyloid-β accumulation and tau spread.

We also report additional exploratory analyses that seek to extend this work by incorporating a more advanced measure of microstructure: hindrance modulated orientational anisotropy (HMOA; Dell’Acqua et al., 2013). HMOA is regarded as a tract-specific measure of microstructure and is argued to be more sensitive to alterations in anisotropy than either FA or MD (Dell’Acqua et al., 2013). As such, we investigated whether *APOE* ε4 is associated with differences in PHCB and ILF HMOA, complementing the primary (replication) analyses. In addition, we also assessed whether *APOE* ε4 is associated with asymmetry in PHCB and ILF HMOA. Recent evidence suggests that AD is characterised by a loss of typical or “healthy” leftward structural and functional asymmetry in the brain (Banks et al., 2018; Roe et al., 2021; Tyrer et al., 2020), perhaps as a result of hemispheric differences in susceptibility to AD pathology (Lubben et al., 2021; Weise et al., 2018). Given the proposal that early-life *APOE* ε4-related alterations in neural activity and connectivity increase vulnerability to AD pathology, notably amyloid-β accumulation but perhaps also tau spread, it is plausible that this allele may be associated with changes in the asymmetry of key white matter tracts. To our knowledge, no study to date has yet investigated this possibility, especially in healthy young adults.

## 2. Method

### 2.1. Participants

Participant data were acquired from a repository at the Cardiff University Brain Research Imaging Centre. Portions of this data have been published elsewhere (Foley et al., 2017; Koelewijn et al., 2019). Participants were healthy adults, who were screened via interview or questionnaire for the presence of neuropsychiatric disorders. All were right-handed, had normal or corrected-to-normal vision, and provided informed consent for their data to be used in imaging genetics analyses. All procedures were originally reviewed and approved by the Cardiff University School of Psychology Research Ethics Committee. For the current study, participants were only included if they completed the requisite MRI scans, had *APOE* genotype information available, and were aged 35 years or under (*N* = 148). After additional exclusions were applied – described below (see also Supplementary Figure 1) – the final sample comprised 128 participants (86 females, 42 males) aged between 19 and 33 years (*M* = 23.8, *SD* = 3.6).

Consistent with Hodgetts et al. (2019), the final sample was split into carrier and non-carrier groups based on the presence/absence of the *APOE* ε4 allele (Table 1). Participants carrying both risk-enhancing (ε4) and risk-reducing (ε2) *APOE* alleles were included as part of the carrier group, as the ε2ε4 genotype is associated with higher levels of AD pathology and risk (Goldberg et al., 2020; Jansen et al., 2015; Reiman et al., 2020). Although *APOE* is often directly genotyped, as in Hodgetts et al.’s study, here it was inferred from imputed (1000G phase 1, version 3) genome-wide genetic data (for more detail, see Foley et al., 2017). Previous research has demonstrated that it is possible to accurately infer *APOE* genotypes using this method (Lupton et al., 2018; Oldmeadow et al., 2014; Radmanesh et al., 2014). Overall, the current sample included 40 *APOE* ε4 carriers (4 ε2/ε4, 33 ε3/ε4, 3 ε4/ε4) and 88 *APOE* ε4 non-carriers (4 ε2/ε2, 14 ε2/ε3, 70 ε3/ε3). An effect size sensitivity analysis calculated using the *pwr* package (version 1.2-2; Champely, 2018) in R (version 3.6.0; R Core Team, 2019) using RStudio (version 1.3.1093; RStudio Team, 2020) revealed that the smallest effect size detectable at 80% power was Cohen’s *d_s_* = 0.575 (1-β = .80, Bonferroni-corrected α = .016, directional hypothesis). By comparison, even without correcting the α level for multiple comparisons, the smallest effect size detectable at 80% power in Hodgetts et al.’s study was Cohen’s *d_s_* = 0.931 (1-β = .80, α = .05, directional hypothesis). Basic sample characteristics in this study and in Hodgetts et al.’s study are compared in Supplementary Table 1.

**Table 1.** Basic Sample Characteristics Separated by APOE ε4 Carrier Status.

|  | APOE ε4+ | APOE ε4- | Statistics |
| --- | --- | --- | --- |
|  | (*n* = 40) | (*n* = 88) |  |
| Age (years; *M* ± *SD*) | 23.9 ± 3.3 | 23.7 ± 3.7 | $t(84.84) = 0.226, p = .822$, Cohen's $d_s = 0.042$, $BF_{10} = 0.206$ |
| Sex (Males/Females; *n*)<sup>a</sup> | 12/28 | 30/58 | $X^2(1, N = 128) = 0.209, p = .648, \phi = 0.04$, $BF_{10} = 0.241$ |

*Note.* Frequentist null hypothesis significance tests (two-sided Welch's *t*-test for age, chi-square test for sex) revealed no significant difference between APOE ε4 carriers and non-carriers in terms of age or sex. Effect sizes were also small, while complementary BF analyses provided moderate evidence in support of the null hypothesis of no difference. Abbreviations: APOE ε4+ = APOE ε4 carrier, APOE ε4- = APOE ε4 non-carrier, *M* = mean, *n* = number of participants, *SD* = standard deviation.
<sup>a</sup>Although sex was self-reported, it was checked against chromosomal sex as part of genetic quality control procedures (Foley et al., 2017).

### 2.2. MRI scan parameters

Scanning was conducted on a GE SIGNA HDx 3T MRI system (General Electric Healthcare, Milwaukee, WI) with an eight-channel receive-only head coil. Whole-brain high angular resolution diffusion imaging data (Tuch et al., 2002) were acquired using a diffusion-weighted single-shot echo-planar imaging sequence (TE = 89ms; voxel dimensions = 2.4 x 2.4 x 2.4mm; FOV = 230mm x 230mm; acquisition matrix = 96 x 96; 60 slices aligned AC/PC with 2.4mm thickness and no gap). Gradients were applied along 30 isotropic directions (Jones et al., 1999) with b = 1200 s/mm^2^. Three non-diffusion-weighted images were acquired with b = 0 s/mm^2^. Acquisitions were cardiac-gated using a peripheral pulse oximeter. T1-weighted anatomical images were acquired using a three-dimensional fast spoiled gradient-echo sequence (TR/TE = 7.8/3s; voxel dimensions = 1mm isotropic; FOV ranging from 256 x 256 x 168mm to 256 x 256 x 180mm; acquisition matrix ranging from 256 x 256 x 168 to 256 x 256 x 180; flip angle = 20°). These sequences were similar to those used by Hodgetts et al. (2019), with only subtle differences between the two studies (outlined in Supplementary Table 2).

### 2.3. dMRI

#### 2.3.1. Pre-processing

The dMRI data were corrected for motion- and eddy current-induced distortions in ExploreDTI (version 4.8.6; Leemans et al., 2009), with an appropriate reorientation of the b-matrix (Leemans & Jones, 2009). Images were registered to down-sampled T1-weighted images (1.5mm isotropic resolution) to correct for susceptibility deformations (Irfanoglu et al., 2012). Data were visually checked as part of quality assurance procedures, leading to the removal of two participants from the analysis due to poor quality data. Consistent with Hodgetts et al. (2019), the two-compartment free-water elimination procedure was implemented using in-house MATLAB code (version R2015a; MathWorks, Inc., 2015) to correct for voxel-wise partial volume artefacts (Pasternak et al., 2009). This procedure has been shown to improve tract delineation, as well as the sensitivity and specificity of measures traditionally derived from the diffusion tensor (Pasternak et al., 2009). Free-water corrected FA and MD maps were then used in further analyses. FA represents the degree to which diffusion is constrained in a particular direction, ranging from 0 (isotropic diffusion) to 1 (anisotropic diffusion). By contrast, MD (10^-3^mm^2^s^-1^) represents the average diffusivity rate.

#### 2.3.2. Tractography

The RESDORE algorithm was used to identify outliers in the diffusion data (Parker, 2014), and then tractography was conducted in ExploreDTI using the modified damped Richardson Lucy spherical deconvolution algorithm (Dell’Acqua et al., 2010). Spherical deconvolution approaches enable multiple peaks to be extracted in the white matter fibre orientation density function (fODF) within a given voxel. This allows complex fibre arrangements, such as crossing/kissing fibres, to be modelled more accurately (Dell’Acqua & Tournier, 2019). The current study and the original study by Hodgetts et al. (2019) both used spherical deconvolution approaches, although the latter used the constrained spherical deconvolution algorithm (Jeurissen et al., 2011). While this might lead to subtle differences between the two studies, the modified damped Richardson Lucy deconvolution algorithm was selected here because it is considered less sensitive to miscalibration (Parker et al., 2013). To minimise any further discrepancies between the studies, tracts were reconstructed using the same parameters used by Hodgetts et al. (fODF amplitude threshold = 0.1; step size = 0.5mm; angle threshold = 60°).

In-house semi-automated tractography software (Parker et al., 2012) was used to generate three-dimensional reconstructions of the PHCB and ILF in both hemispheres. The software was trained on manual reconstructions generated by author R.L. using a waypoint ROI approach in ExploreDTI, where “SEED”, “AND”, and “NOT” ROIs were used to isolate tract-specific streamlines (Figure 1). ROIs were placed in the same regions as described by Hodgetts et al. (2019). Placement was therefore guided by established protocols for the PHCB (Jones, Christiansen et al., 2013) and the ILF (Wakana et al., 2007), respectively. All reconstructions generated by the semi-automated software were visually inspected by authors R.L. and C.J.H. and, where required, manually edited post hoc to remove erroneous, anatomically implausible fibres. Participants for whom the PHCB and ILF could not be reconstructed in both hemispheres were removed from analysis (*n* = 18). Thereafter, measures of microstructure were obtained and averaged across tracts. Although the semi-automated approach used here differs to that used by Hodgetts et al., larger studies have shown this to be useful (Foley et al., 2017; Metzler-Baddeley et al., 2019). Furthermore, during visual inspection, author C.J.H. confirmed that tract reconstruction produced qualitatively similar outputs to those obtained in the original, to-be-replicated study.

**Figure 1.**
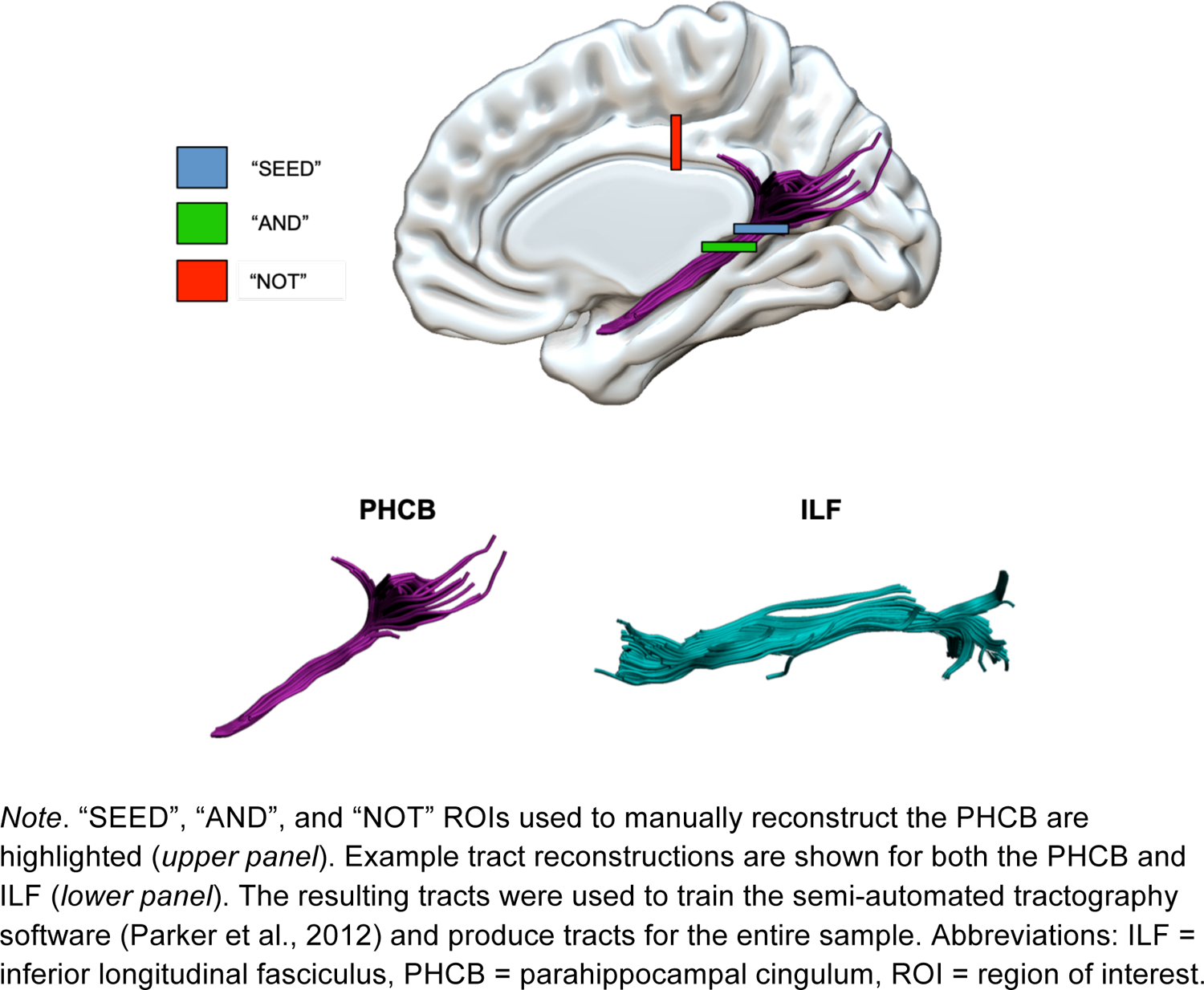
Manual Reconstructions of the PHCB and ILF

#### 2.3.3. Tract-based spatial statistics (TBSS)

Complementary voxel-wise statistical analysis of the FA and MD data was conducted using TBSS (Smith et al., 2006). Each participant’s free-water corrected FA and MD maps were first aligned in standard MNI space using nonlinear registration (Andersson et al., 2007a, 2007b). Next, the mean FA images were created and subsequently thinned (threshold = 0.2) to generate the mean FA skeleton, which represents the centre of all tracts common to the group. Each participant’s aligned FA and MD data were then projected onto the skeleton and the resulting data carried forward for voxel-wise cross-subject analysis. These analyses were performed using *randomise* (Winkler et al., 2014), a permutation-based inference tool. For both FA and MD, a general linear model contrasting *APOE* ε4 carriers and non-carriers (FA: carrier > non-carrier; MD: carrier < non-carrier) was applied (*n* permutations = 1000). Mirroring Hodgetts et al.’s (2019) example, analyses were first restricted to the PHCB using an ROI mask [labelled “cingulum (hippo-campus)”] from the John Hopkins University ICBM-DTI-81 white-matter tractography atlas. An exploratory whole-brain analysis was then conducted. Statistically significant clusters were extracted from both analyses using threshold-free cluster enhancement with a corrected α level of 0.05 (Smith and Nichols, 2009).

### 2.4. Statistical analyses

Except for TBSS, all statistical analyses were conducted using R in RStudio. In addition to common frequentist null hypothesis significance tests, Bayes factors (BFs) were calculated. BFs quantify the degree to which the observed data favours predictions made by two models, in this case the null hypothesis and the alternative hypothesis. Consequently, BF analyses can provide evidence in support of the null (Dienes, 2014). In accordance with the evidence categories outlined by Lee and Wagenmakers (2013), a BF_+0_ (BF_10_ for two-sided tests) greater than 3 was considered to represent at least moderate evidence for the alternative hypothesis, whereas a BF_+0_ less than .33 was considered to represent at least moderate evidence for the null hypothesis.

#### 2.4.1. Primary (replication) analyses

To test whether *APOE* ε4 carriers showed higher FA and lower MD in the PHCB but not the ILF, one-sided Welch’s *t*-tests were conducted. As in Hodgetts et al. (2019), all tests were repeated, once with male participants removed and once with ε2 carriers removed. These additional tests – performed independently of each other – were originally conducted based on evidence that *APOE* ε4 may have a stronger effect on AD biomarkers in females than males (Riedel et al., 2016), whereas *APOE* ε2 may have a protective effect on AD biomarkers (Suri et al., 2013). To ensure that the probability of falsely rejecting the null – the Type I error rate – was not inflated, a Bonferroni correction was applied to the α level (.05 / 3 = .016). Two BFs were also calculated: a default JZS BF and a replication BF. The default JZS BF, which uses a default prior distribution and was computed using the *BayesFactor* package (version 0.9.12-4.2; Morey & Rouder, 2018), examines whether an effect is present or absent in the data collected in the replication study regardless of the original effect. Here, one-sided (directional) default JZS BFs were calculated. The replication BF, by contrast, uses the posterior distribution of the original study as the prior distribution in the replication study, examining whether the original effect is present or absent in the data collected in the replication study. This BF was computed using previously published R code (Verhagen & Wagenmakers, 2014).

#### 2.4.2. Secondary (extension) analyses

##### 2.4.2.1. HMOA index

It remains to be seen whether *APOE* ε4-related differences in PHCB microstructure are better captured by measures other than FA and MD, which are sensitive to various aspects of white matter microstructure without being specific to any one (Jones, Knösche, & Turner, 2013). One such measure is HMOA, which is defined as the absolute amplitude of each fODF lobe (Dell’Acqua et al., 2013). This is normalised using a reference amplitude in order to create an index bound between zero and one. A value of zero reflects the absence of a fibre, whereas a value of one reflects the highest fODF signal that can realistically be detected in biological tissue (Dell’Acqua et al., 2013).

Given the lack of a directional hypothesis relating to HMOA, two-sided Welch’s *t*-tests and two-sided default JZS BFs were used to identify any differences between *APOE* ε4 carriers and non-carriers. In keeping with the primary (replication) analyses described above, these tests were repeated with males removed and then with ε2 carriers removed. These analytical steps were performed independently. A Bonferroni correction was applied to the nominal α level (.05 / 3 = .016).

##### 2.4.2.2. Hemispheric asymmetry

Despite reports linking AD with a loss of leftward structural and functional asymmetry (Banks et al., 2018; Roe et al., 2021; Tyrer et al., 2020), which may be related to differences in hemispheric susceptibility to pathology (Lubben et al., 2021; Weise et al., 2018), no study to our knowledge has yet investigated whether the *APOE* ε4 allele is associated with asymmetry in PHCB or ILF microstructure. Moreover, considering the proposed interaction between *APOE* ε4 and sex in the context of AD risk (Riedel et al., 2016), there is also an interesting question as to whether sex moderates any potential *APOE* ε4-related association with hemispheric asymmetry. We therefore examined whether HMOA – a more tract-specific measure – was lateralised to the left or right hemisphere, and whether this was impacted by *APOE* ε4, sex, or their interaction.

As with the analyses described previously, the ILF was included as a comparison tract. Lateralisation indices (LIs) were calculated for HMOA in both the PHCB and ILF [LI = (right - left) / (right + left)]. For any given participant, a negative LI score indicates that HMOA was higher in the left hemisphere, whereas a positive LI score indicates that HMOA was higher in the right hemisphere (Zhao et al., 2016). These LI_HMOA_ scores were subsequently analysed using robust multiple linear regression, which was carried out via the *lmrob* function from the *robustbase* package (version 0.93-7; Maechler et al., 2021). The fitted models were as follows:

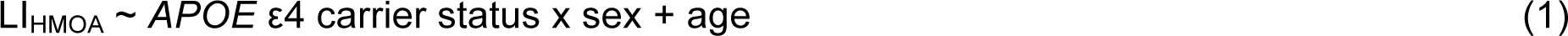

LIs were entered as dependent variables. *APOE* ε4 carrier status and sex were treated as categorical variables and coded using deviation coding. Age – included as a covariate of “no interest” – was centred and scaled. The interaction between *APOE* ε4 carrier status and sex was included in the model. Results were deemed statistically significant if the observed *p* value was smaller than the nominal α level of 0.05.

### 2.5. Data and code availability

R code used to analyse and visualise data in the current study is made publicly available via the Open Science Framework (https://osf.io/f6jp3/). Due to the sensitive nature of the data, the original ethics do not allow for the public archiving of study data (for more information, see Koelewijn et al., 2019). Access to pseudo-anonymised data may be granted, however, after the signing and approval of suitable data-transfer agreements. Readers seeking access through this mechanism should contact Professor Krish D. Singh at the Cardiff University Brain Research Imaging Centre.

## 3. Results

### 3.1. Primary (replication) analyses

#### 3.1.1. Effect of *APOE* ε4 on PHCB FA and MD

FA values for the PHCB – separated by *APOE* ε4 carrier status – are shown in Figure 2A. Contrary to our initial hypothesis, PHCB FA was not significantly higher for *APOE* ε4 carriers than non-carriers (*t*(87.559) = −0.606, *p* = .727, Cohen’s *d_s_* = −0.112). Supporting this, BF analysis produced moderate evidence in favour of the null (default JZS BF_+0_ = 0.138, replication BF_10_ = 0.141). Removing males from the analysis did not alter the results in any meaningful way (*t*(57.685) = 0.045, *p* = .482, Cohen’s *d_s_* = 0.01, default JZS BF_+0_ = 0.246, replication BF_10_ = 0.168), nor did removing ε2 carriers (*t*(84.459) = −0.923, *p* = .821, Cohen’s *d_s_* = −0.183, default JZS BF_+0_ = 0.125, replication BF_10_ = 0.271).

**Figure 2.**
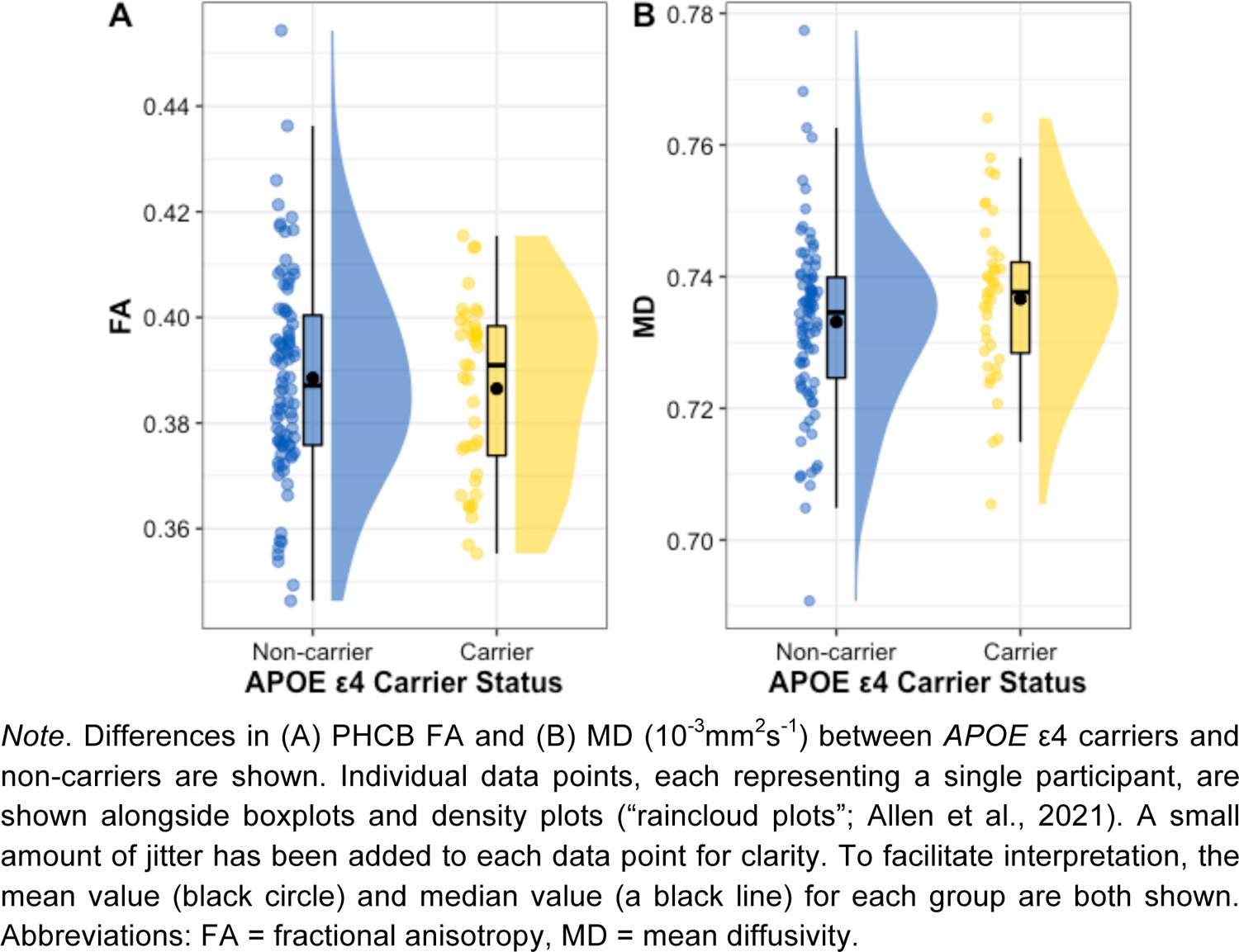
Differences in PHCB FA and MD Between APOE ε4 Carriers and Non-Carriers

MD values for the PHCB – separated by *APOE* ε4 carrier status – are shown in Figure 2B. Again, contrary to prior expectations, PHCB MD was not significantly lower for *APOE* ε4 carriers than non-carriers (*t*(83.625) = 1.429, *p* = .922, Cohen’s *d_s_* = 0.267). Here, BF analysis revealed strong evidence in favour of the null (default JZS BF_+0_ = 0.092, replication BF_10_ = 0.057). As with FA, the results for MD did not change substantively after removing males (*t*(59.729) = 1.515, *p* = .933, Cohen’s *d_s_* = 0.341, default JZS BF_+0_ = 0.106, replication BF_10_ = 0.054) or after removing ε2 carriers (*t*(79.581) = 1.328, *p* = .906, Cohen’s *d_s_* = 0.267, default JZS BF_+0_ = 0.103, replication BF_10_ = 0.1).

#### 3.1.2. Effect of *APOE* ε4 on ILF FA and MD

The same analysis was conducted on ILF FA and MD. Analysis revealed that ILF FA was not significantly higher for *APOE* ε4 carriers than non-carriers (*t*(86.143) = −0.864, *p* = .805, Cohen’s *d_s_* = −0.16). BF analysis provided moderate-to-strong evidence favouring the absence of an effect (default JZS BF_+0_ = 0.12), as well as anecdotal-to-moderate evidence favouring the absence of the effect reported by Hodgetts et al. (replication BF_10_ = 0.309). This slight discrepancy between BFs is likely because the original to-be-replicated effect was also small and did not reach the threshold for statistical significance, meaning that the informed prior used was already more “sceptical” than the default prior. Results remained largely unchanged when males were removed (*t*(49.129) = −0.069, *p* = .527, Cohen’s *d_s_* = − 0.016, default JZS BF_+0_ = 0.226, replication BF_10_ = 0.308) and when ε2 carriers were removed (*t*(79.5) = −0.893, *p* = .813, Cohen’s *d_s_* = −0.179, default JZS BF_+0_ = 0.126).

ILF MD was not significantly lower for *APOE* ε4 carriers than non-carriers (*t*(81.941) = 0.54, *p* = .705, Cohen’s *d_s_* = 0.101). BFs again provided evidence in support of the null (default JZS BF_+0_ = 0.142, replication BF_10_ = 0.446). Removing males had no notable impact on the results (*t*(55.856) = 0.818, *p* = .792, Cohen’s *d_s_* = 0.187, default JZS BF_+0_ = 0.144, replication BF_10_ = 0.613) nor did removing *APOE* ε2 carriers (*t*(75.242) = 0.713, *p* = .761, Cohen’s *d_s_* = 0.145, default JZS BF_+0_ = 0.137).

#### 3.1.2. TBSS

Consistent with the tractography analysis, PHCB-restricted TBSS analysis revealed no significant differences between *APOE* ε4 carriers and non-carriers. This was true of both FA (contrast: carriers > non-carriers) and MD (contrast: carriers < non-carriers). Adopting an uncorrected α level of *p* = .005, as has been done previously (Hodgetts et al., 2019; Postans et al., 2014), did not alter this outcome. Exploratory whole-brain TBSS analysis provided complementary evidence, with no differences evident between *APOE* ε4 carriers and non-carriers.

### 3.2. Secondary (extension) analyses

#### 3.2.1. Effect of APOE ε4 on PHCB and ILF HMOA

Analysis revealed no significant difference between *APOE* ε4 carriers and non-carriers in terms of PHCB HMOA (*t*(90.357) = −0.399, *p* = .691, Cohen’s *d_s_* = −0.073). BF analysis also provided moderate evidence in favour of the null (default JZS BF_10_ = 0.215). These results were largely unaffected by the removal of males (*t*(58.33) = 0.445, *p* = .658, Cohen’s *d_s_* = 0.10, default JZS BF_10_ = 0.258) or the removal of ε2 carriers (*t*(85.926) = −0.844, *p* = .401, Cohen’s *d_s_* = −0.167, default JZS BF_10_ = 0.283).

For completeness, the same analysis was conducted for ILF HMOA. Results revealed that *APOE* ε4 carriers and non-carriers did not differ significantly in terms of ILF HMOA (*t*(94.682) = −0.762, *p* = .448, Cohen’s *d_s_* = −0.139). BF analysis provided complementary evidence, largely favouring the null (default JZS BF_10_ = 0.251). This remained the case when males were removed (*t*(48.941) = 0.394, *p* = .696, Cohen’s *d_s_* = 0.092, default JZS BF_10_ = 0.256) and when individuals possessing the ε2 allele were removed (*t*(84.914) = −0.819, *p* = .415, Cohen’s *d_s_* = −0.162, default JZS BF_10_ = 0.279).

#### 3.2.2. Hemispheric asymmetry in PHCB and ILF HMOA

In terms of hemispheric asymmetry, analysis revealed that HMOA was higher in the right (*M* = .234, *SD* = .015) than the left (*M* = .224, *SD* = .018) PHCB (*t*(127) = −6.631, *p* < .001, Cohen’s *d_z_* = −0.586, default JZS BF_10_ > 100). Nevertheless, for PHCB LI_HMOA_, there was no significant association with *APOE* ε4 (*b* < −.001, *p* = .911), sex (*b* = −.002, *p* = .743), or their interaction (*b* = −.008, *p* = .558). Consequently, we observed no evidence indicating that *APOE* ε4, sex, or their interactions influenced hemispheric asymmetry in PHCB microstructure.

The same analysis was conducted on ILF microstructure. HMOA was higher in the left (*M* = .302, *SD* = .027) than the right (*M* = .293, *SD* = .029) hemisphere (*t*(127) = 3.778, *p* < .001, Cohen’s *d_z_* = 0.334, default JZS BF_10_ = 74.09). Examining whether this hemispheric asymmetry was influenced by *APOE* ε4, sex, or their interaction, LIs were again calculated and analysed. In the case of ILF LI_HMOA_, there was a significant association with *APOE* ε4 (*b* = 0.027, *p* = .005) but not with sex (*b* = 0.014, *p* = .156) or their interaction (*b* = 0.007, *p* = .674). Figure 3 highlights the group-level differences in ILF LI_HMOA_. As shown, this was driven by reduced leftward asymmetry in this tract among *APOE* ε4 carriers than non-carriers.

**Figure 3.**
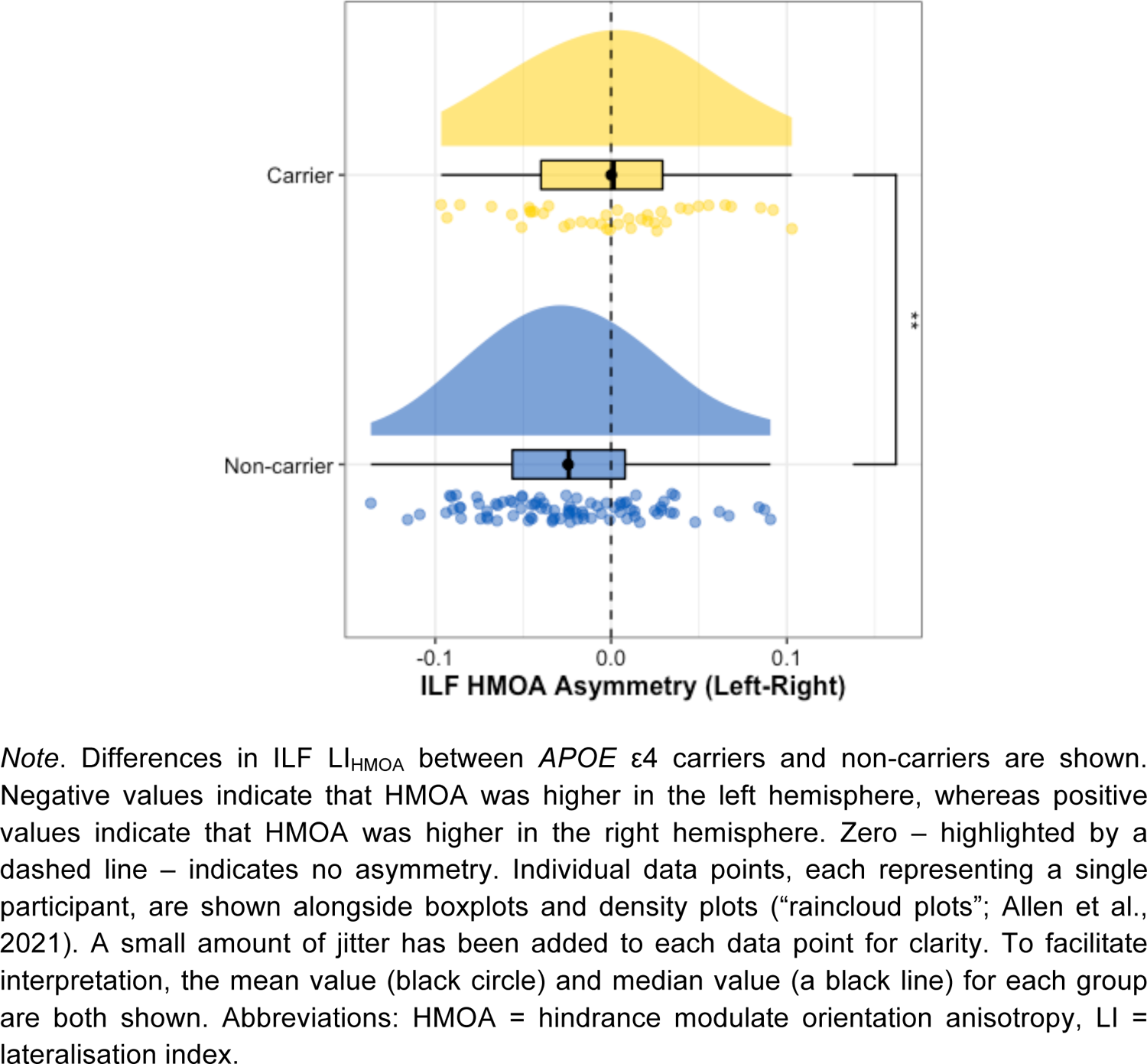
Difference in ILF LI_HMOA_ Between APOE ε4 Carriers and Non-Carriers

## 4. Discussion

In this study, we aimed to replicate Hodgetts et al.’s (2019) findings that healthy young *APOE* ε4 carriers show higher FA and lower MD than non-carriers in the PHCB but not the ILF. Such a pattern would be in line with suggestions that individuals with pre-existing “hyper-connectivity” between posteromedial cortex and the medial temporal lobe may be more vulnerable to amyloid-β accumulation (Buckner et al., 2009; Bero et al., 2012; de Haan et al., 2012) and/or tau spread (Jacobs et al., 2018; Ziontz et al., 2021). Extending this work, we also conducted analyses on HMOA, a measure that is proposed to be more sensitive to alterations in tract microstructure than FA or MD (Dell’Acqua et al., 2013). This included an investigation into hemispheric asymmetry in PHCB and ILF HMOA, as prior reports indicate that AD impacts brain asymmetry (Banks et al., 2018; Roe et al., 2021; Tyrer et al., 2020).

In contrast to the original study, we did not observe higher FA or lower MD in the PHCB of young *APOE* ε4 carriers compared to non-carriers. Rather, we found: no statistically significant effects in the expected direction (all *p*s ≥ .482); relatively small effect sizes (Cohen’s *d_s_* range from −0.183 to 0.341); and BFs providing evidence in favour of the null (default JZS BF_+0_ range from .092 to .246, replication BF_10_ range from .054 to .273). Crucially, these BFs represent moderate-to-strong evidence in support of the null hypothesis (Lee & Wagenmakers, 2013). As such, we not only failed to replicate the effect reported by Hodgetts et al. (2019), but also found evidence against the presence of such an effect. There are several plausible explanations for this, although they are not necessarily mutually exclusive.

First, it could be the case that Hodgetts et al.’s (2019) findings were false positives (see also Dell’Acqua et al., 2015). Hodgetts et al.’s study included just 15 participants in the *APOE* ε4 carrier and non-carrier groups and, as such, was likely underpowered to detect an effect of the magnitude one might expect from this common genetic variant, especially in early adulthood (Henson et al., 2020). Given that low statistical power reduces the probability that an observed effect represents a true effect (Button et al., 2013), it is possible that the effects reported by Hodgetts et al. were false positives, although it is unclear how this relates the their observation that PHCB microstructure correlated with posteromedial cortex activity during perceptual scene discrimination (see also Shine et al., 2015). The BF analyses conducted here provide complementary support for this assertion, demonstrating that the observed data favour the null. Taken at face value, this interpretation casts doubt on the notion that increased connectivity between posteromedial cortex and the medial temporal lobe – mediated by individual differences in PHCB microstructure – represents a pre-existing *APOE* ε4-related trait enhancing vulnerability to amyloid-β accumulation and/or tau spread.

Alternatively, it could be the case that Hodgetts et al. (2019) observed a true effect, but its magnitude was exaggerated. Effect size inflation is most likely to occur in studies with small sample sizes, a phenomenon referred to as the “winner’s curse” (Button et al., 2013). If true, the analysis reported in this replication attempt might itself be underpowered to detect the effect of *APOE* ε4 on PHCB FA and MD, thereby constituting a Type II error or false negative. Such an explanation would help to reconcile the observed findings with prior results indicating that *APOE* ε4 does have an impact on posteromedial connectivity early in life (Brown et al., 2011; Felsky & Voineskos, 2013; Hodgetts et al., 2019). While this cannot currently be ruled out, it should be noted that an effect size sensitivity analysis revealed that the smallest effect size detectable at 80% power in the current study was Cohen’s *d_s_* = 0.57. In addition, the BF analyses conducted here indicated that the observed data provided moderate-to-strong evidence in favour of the null, as opposed to simply providing inconclusive evidence. This shows that, even with the current sample size, our findings have relatively high evidential value (Dienes, 2014).

Another potential explanation is that the *APOE* ε4 carriers and non-carriers included in the two studies differed in other AD-relevant factors. It is well established that while *APOE* ε4 carriers are at increased risk of developing AD relative to non-carriers, not all go on to develop the disease (Liu et al., 2013). In fact, only ∼50% of individuals with AD possess one or more copies of the *APOE* ε4 allele (Karch et al., 2014), highlighting the importance of other factors – genetic and environmental – in disease risk/protection (Jagust & Mormino, 2011; Silva et al., 2019). Following this line of reasoning, it is possible that – due to sampling variation – the *APOE* ε4 carrier and non-carrier groups included in the two studies differed in their overall AD risk profiles, with potential implications for white matter microstructure. This would at least partly explain why we failed to replicate the effect originally reported by Hodgetts et al. (2019). Nevertheless, it is important to recognise that this remains an open question, and large-scale dMRI studies are required to test this possibility.

Regarding the asymmetry of PHCB microstructure, we found that HMOA was higher in the right hemisphere. This is consistent with some previous reports using diffusion tensor metrics (Metzler-Baddeley et al., 2012; Powell et al., 2012), although certainly not all (Lebel et al., 2012; Thiebaut de Schotten et al., 2011). Prior research has suggested that while left-hemispheric networks exhibit increased nodal efficiency in brains areas supporting language, right-hemispheric networks exhibit increased nodal efficiency in brain areas related to episodic memory (Caeyenberghs & Leemans, 2014). This potentially highlights a functional role for the observed rightward asymmetry in PHCB microstructure. However, we did not observe an effect of *APOE* ε4 or sex on the degree of PHCB asymmetry.

A different pattern emerged in the analysis of ILF microstructure, with HMOA characterised by leftward asymmetry. As with the PHCB, this finding is consistent with a number of studies examining asymmetry in ILF volume and diffusion tensor-derived measures of microstructure (Banfi et al., 2019; Panesar et al., 2018; Thiebaut de Schotten et al., 2011). We also observed that the degree of asymmetry in this tract was associated with *APOE* ε4 carrier status, such that asymmetry was lower in carriers relative to non-carriers, mirroring to some extent the loss of leftward asymmetry in AD (Banks et al., 2018; Roe et al., 2021; Tyrer et al., 2020). The ILF connects occipital and ventro-anterior temporal lobe (Herbet et al., 2018), underpinning a network involved in representing item information, including semantic and perceptual information (Murray et al., 2017; Ranganath & Ritchey, 2012).

Recent research suggests that complex item discrimination is impaired in AD risk (Fidalgo et al., 2016; Mason et al., 2017), which has in turn been linked to the structure and function of components within this network (Berron et al., 2018; Olsen et al., 2017; Reagh et al., 2016). Indeed, complex item discrimination has been proposed as a useful measure for the detection of early AD (Gaynor et al., 2019). In addition, a recent study of young adult *APOE* ε4 carriers in the Human Connectome Project failed to replicate enhanced intrinsic functional connectivity between posteromedial cortex and the medial temporal lobe, as observed previously (Filippini et al., 2009), but found heightened activity in left hemisphere regions connected by the ILF during face encoding (Mentink et al., 2021), possibly suggestive of a lifelong neural inefficiency (Jagust & Mormino, 2011). Future research should seek to replicate further the effect of *APOE* ε4 on reduced structural (and functional) left hemispheric asymmetry, especially given potential implications for later life cognition (Jiang et al., 2021; Maass et al., 2019).

## 5. Summary

In this study, we failed to replicate Hodgetts et al.’s (2019) finding that, relative to non-carriers, healthy young adult *APOE* ε4 carriers show higher FA and lower MD in the PHCB but not the ILF. Rather, the observed data strongly supported the null hypothesis of no difference. Our findings thus suggest that young adult *APOE* ε4 carriers do not show alterations in PHCB microstructure that might enhance vulnerability – via excessive connectivity-dependent neuronal activity – to amyloid-β accumulation and/or tau spread.

Nevertheless, marked patterns of hemispheric asymmetry were evident in PHCB and ILF microstructure, although only the latter was associated with *APOE* ε4 carrier status. Given the potential implications for later life cognition, our study highlights an important area for future research seeking to understand how this AD risk factor impacts neural and cognitive efficiency years prior to the onset of clinical symptoms.

## Acknowledgements

We would like to thank Ofer Pasternak for providing the free-water elimination pipeline, Sonya Foley for assistance in identifying relevant data in the repository, and Mark Postans for helpful discussions.

## References

1. Acosta-Cabronero, J., Williams, G. B., Pengas, G., & Nestor, P. J. (2010). Absolute diffusivities define the landscape of white matter degeneration in Alzheimer’s disease. Brain, 133(2), 529–539. https://doi.org/10.1093/brain/awp257

2. Albert, M. S., DeKosky, S. T., Dickson, D., Dubois, B., Feldman, H. H., Fox, N. C., Gamst, A., Holtzman, D. M., Jagust, W. J., Petersen, R. C., Snyder, P. J., Carrillo, M. C., Thies, B., & Phelps, C. H. (2011). The diagnosis of mild cognitive impairment due to Alzheimer’s disease: Recommendations from the National Institute on Aging-Alzheimer’s Association workgroups on diagnostic guidelines for Alzheimer’s disease. Alzheimer’s & Dementia, 7(3), 270–279. https://doi.org/10.1016/j.jalz.2011.03.008

3. Allen, M., Poggiali, D., Whitaker, K., Marshall, T. R., van Langen, J., & Kievit, R. A. (2021). Raincloud plots: A multi-platform tool for robust data visualization. Wellcome Open Research, 4, 63. https://doi.org/10.12688/wellcomeopenres.15191.2

4. Andersson, J. L. R., Jenkinson, M., & Smith, S. (2007a). Non-linear optimisation. FMRIB technical report TR07JA1. www.fmrib.ox.ac.uk/analysis/techrep

5. Andersson, J. L. R., Jenkinson, M., & Smith, S. (2007b). Non-linear registration, aka spatial normalisation FMRIB technical report TR07JA2. www.fmrib.ox.ac.uk/analysis/techrep

6. Assaf, Y., Johansen-Berg, H., & Thiebaut de Schotten, M. (2019). The role of diffusion MRI in neuroscience. NMR in Biomedicine, 32(4), e3762. https://doi.org/10.1002/nbm.3762

7. Banfi, C., Koschutnig, K., Moll, K., Schulte-Körne, G., Fink, A., & Landerl, K. (2019). White matter alterations and tract lateralization in children with dyslexia and isolated spelling deficits. Human Brain Mapping, 40(3), 765–776. https://doi.org/10.1002/hbm.24410

8. Banks, S. J., Zhuang, X., Bayram, E., Bird, C., Cordes, D., Caldwell, J. Z. K., Cummings, J. L., & for the Alzheimer’s Disease Neuroimaging Initiative. (2018). Default mode network lateralization and memory in healthy aging and Alzheimer’s disease. Journal of Alzheimer’s Disease, 66(3), 1223–1234. https://doi.org/10.3233/JAD-180541

9. Belloy, M. E., Napolioni, V., & Greicius, M. D. (2019). A quarter century of APOE and Alzheimer’s disease: Progress to date and the path forward. Neuron, 101(5), 820–838. https://doi.org/10.1016/j.neuron.2019.01.056

10. Benitez, A., Jensen, J. H., Falangola, M. F., Spampinato, M. V., Rieter, W. J., Nietert, P. J., Fountain-Zaragoza, S., Keith, K., Dhiman, S., & Helpern, J. A. (2021). Greater diffusion restriction in white matter tracts in preclinical AD. Alzheimer’s & Dementia, 17(S5), e054942. https://doi.org/10.1002/alz.054942

11. Bero, A. W., Bauer, A. Q., Stewart, F. R., White, B. R., Cirrito, J. R., Raichle, M. E., Culver, J. P., & Holtzman, D. M. (2012). Bidirectional relationship between functional connectivity and amyloid-β deposition in mouse brain. Journal of Neuroscience, 32(13), 4334–4340. https://doi.org/10.1523/JNEUROSCI.5845-11.2012

12. Berron, D., Neumann, K., Maass, A., Schütze, H., Fliessbach, K., Kiven, V., Jessen, F., Sauvage, M., Kumaran, D., & Düzel, E. (2018). Age-related functional changes in domain-specific medial temporal lobe pathways. Neurobiology of Aging, 65, 86–97. https://doi.org/10.1016/j.neurobiolaging.2017.12.030

13. Bozzali, M., Giulietti, G., Basile, B., Serra, L., Spanò, B., Perri, R., Giubilei, F., Marra, C., Caltagirone, C., & Cercignani, M. (2012). Damage to the cingulum contributes to Alzheimer’s disease pathophysiology by deafferentation mechanism. Human Brain Mapping, 33(6), 1295–1308. https://doi.org/10.1002/hbm.21287

14. Brown, J. A., Terashima, K. H., Burggren, A. C., Ercoli, L. M., Miller, K. J., Small, G. W., & Bookheimer, S. Y. (2011). Brain network local interconnectivity loss in aging APOE-4 allele carriers. Proceedings of the National Academy of Sciences, 108(51), 20760– 20765. https://doi.org/10.1073/pnas.1109038108

15. Bubb, E. J., Metzler-Baddeley, C., & Aggleton, J. P. (2018). The cingulum bundle: Anatomy, function, and dysfunction. Neuroscience & Biobehavioral Reviews, 92, 104–127. https://doi.org/10.1016/j.neubiorev.2018.05.008

16. Buckner, R. L., Sepulcre, J., Talukdar, T., Krienen, F. M., Liu, H., Hedden, T., Andrews-Hanna, J. R., Sperling, R. A., & Johnson, K. A. (2009). Cortical hubs revealed by intrinsic functional connectivity: Mapping, assessment of stability, and relation to Alzheimer’s disease. Journal of Neuroscience, 29(6), 1860–1873. https://doi.org/10.1523/JNEUROSCI.5062-08.2009

17. Burnham, S. C., Laws, S. M., Budgeon, C. A., Doré, V., Porter, T., Bourgeat, P., Buckley, R. F., Murray, K., Ellis, K. A., Turlach, B. A., Salvado, O., Ames, D., Martins, R. N., Rentz, D., Masters, C. L., Rowe, C. C., & Villemagne, V. L. (2020). Impact of APOE-ε4 carriage on the onset and rates of neocortical Aβ-amyloid deposition. Neurobiology of Aging, 95, 46–55. https://doi.org/10.1016/j.neurobiolaging.2020.06.001

18. Button, K. S., Ioannidis, J. P. A., Mokrysz, C., Nosek, B. A., Flint, J., Robinson, E. S. J., & Munafò, M. R. (2013). Power failure: Why small sample size undermines the reliability of neuroscience. Nature Reviews Neuroscience, 14(5), 365–376. https://doi.org/10.1038/nrn3475

19. Caeyenberghs, K., & Leemans, A. (2014). Hemispheric lateralization of topological organization in structural brain networks. Human Brain Mapping, 35(9), 4944–4957. https://doi.org/10.1002/hbm.22524

20. Champely, S. (2018). pwr: Basic functions for power analysis (Version 1.2-2) [Computer software]. https://CRAN.R-project.org/package=pwr

21. Choo, I. H., Lee, D. Y., Oh, J. S., Lee, J. S., Lee, D. S., Song, I. C., Youn, J. C., Kim, S. G., Kim, K. W., Jhoo, J. H., & Woo, J. I. (2010). Posterior cingulate cortex atrophy and regional cingulum disruption in mild cognitive impairment and Alzheimer’s disease. Neurobiology of Aging, 31(5), 772–779. https://doi.org/10.1016/j.neurobiolaging.2008.06.015

22. Collij, L. E., Ingala, S., Top, H., Wottschel, V., Stickney, K. E., Tomassen, J., Konijnenberg, E., ten Kate, M., Sudre, C., Lopes Alves, I., Yaqub, M. M., Wink, A. M., Van‘t Ent, D., Scheltens, P., van Berckel, B. N. M., Visser, P. J., Barkhof, F., & Braber, A. D. (2021). White matter microstructure disruption in early stage amyloid pathology. *Alzheimer’s & Dementia: Diagnosis*, Assessment & Disease Monitoring, 13(1), e12124. https://doi.org/10.1002/dad2.12124

23. Coughlan, G., Laczó, J., Hort, J., Minihane, A.-M., & Hornberger, M. (2018). Spatial navigation deficits—Overlooked cognitive marker for preclinical Alzheimer disease? Nature Reviews Neurology, 14(8), 496–506. https://doi.org/10.1038/s41582-018-0031-x

24. Dalboni da Rocha, J. L., Bramati, I., Coutinho, G., Tovar Moll, F., & Sitaram, R. (2020). Fractional anisotropy changes in parahippocampal cingulum due to Alzheimer’s disease. Scientific Reports, 10, 2660. https://doi.org/10.1038/s41598-020-59327-2

25. de Haan, W., Mott, K., van Straaten, E. C. W., Scheltens, P., & Stam, C. J. (2012). Activity dependent degeneration explains hub vulnerability in Alzheimer’s disease. PLOS Computational Biology, 8(8), e1002582. https://doi.org/10.1371/journal.pcbi.1002582

26. Dell’Acqua, F., Khan, W., Gottlieb, N., Giampietro, V., Ginestet, C., Bouls, D., Newhouse, S., Dobson, R., Banaschewski, T., Barker, G. J., Bokde, A. L. W., Büchel, C., Conrod, P., Flor, H., Frouin, V., Garavan, H., Gowland, P., Heinz, A., Lemaítre, H., … the IMAGEN consortium. (2015). Tract based spatial statistic reveals no differences in white matter microstructural organization between carriers and non-carriers of the APOE ɛ4 and ɛ2 alleles in young healthy adolescents. Journal of Alzheimer’s Disease, 47(4), 977–984. https://doi.org/10.3233/JAD-140519

27. Dell’Acqua, F., Scifo, P., Rizzo, G., Catani, M., Simmons, A., Scotti, G., & Fazio, F. (2010). A modified damped Richardson–Lucy algorithm to reduce isotropic background effects in spherical deconvolution. NeuroImage, 49(2), 1446–1458. https://doi.org/10.1016/j.neuroimage.2009.09.033

28. Dell’Acqua, F., Simmons, A., Williams, S. C. R., & Catani, M. (2013). Can spherical deconvolution provide more information than fiber orientations? Hindrance modulated orientational anisotropy, a true-tract specific index to characterize white matter diffusion. Human Brain Mapping, 34(10), 2464–2483. https://doi.org/10.1002/hbm.22080

29. Dell’Acqua, F., & Tournier, J.-D. (2019). Modelling white matter with spherical deconvolution: How and why? NMR in Biomedicine, 32(4), e3945. https://doi.org/10.1002/nbm.3945

30. DeTure, M. A., & Dickson, D. W. (2019). The neuropathological diagnosis of Alzheimer’s disease. Molecular Neurodegeneration, 14, 32. https://doi.org/10.1186/s13024-019-0333-5

31. Dienes, Z. (2014). Using Bayes to get the most out of non-significant results. Frontiers in Psychology, 5, 781. https://doi.org/10.3389/fpsyg.2014.00781

32. Dong, J. W., Jelescu, I. O., Ades-Aron, B., Novikov, D. S., Friedman, K., Babb, J. S., Osorio, R. S., Galvin, J. E., Shepherd, T. M., & Fieremans, E. (2020). Diffusion MRI biomarkers of white matter microstructure vary nonmonotonically with increasing cerebral amyloid deposition. Neurobiology of Aging, 89, 118–128. https://doi.org/10.1016/j.neurobiolaging.2020.01.009

33. Felsky, D., & Voineskos, A. N. (2013). APOE ɛ4, aging, and effects on white matter across the adult life span. JAMA Psychiatry, 70(6), 646–647. https://doi.org/10.1001/jamapsychiatry.2013.865

34. Fidalgo, C. O., Changoor, A. T., Page-Gould, E., Lee, A. C. H., & Barense, M. D. (2016). Early cognitive decline in older adults better predicts object than scene recognition performance. Hippocampus, 26(12), 1579–1592. https://doi.org/10.1002/hipo.22658

35. Filippini, N., MacIntosh, B. J., Hough, M. G., Goodwin, G. M., Frisoni, G. B., Smith, S. M., Matthews, P. M., Beckmann, C. F., & Mackay, C. E. (2009). Distinct patterns of brain activity in young carriers of the APOE-ε4 allele. Proceedings of the National Academy of Sciences, 106(17), 7209–7214. https://doi.org/10.1073/pnas.0811879106

36. Foley, S. F., Tansey, K. E., Caseras, X., Lancaster, T., Bracht, T., Parker, G., Hall, J., Williams, J., & Linden, D. E. J. (2017). Multimodal brain imaging reveals structural differences in Alzheimer’s disease polygenic risk carriers: A study in healthy young adults. Biological Psychiatry, 81(2), 154–161. https://doi.org/10.1016/j.biopsych.2016.02.033

37. Frisoni, G. B., Altomare, D., Thal, D. R., Ribaldi, F., van der Kant, R., Ossenkoppele, R., Blennow, K., Cummings, J., van Duijn, C., Nilsson, P. M., Dietrich, P.-Y., Scheltens, P., & Dubois, B. (2022). The probabilistic model of Alzheimer disease: The amyloid hypothesis revised. Nature Reviews Neuroscience, 23(1), 53–66. https://doi.org/10.1038/s41583-021-00533-w

38. Gaynor, L. S., Curiel, R. E., Penate, A., Rosselli, M., Burke, S. N., Wicklund, M., Loewenstein, D. A., & Bauer, R. M. (2019). Visual object discrimination impairment as an early predictor of Mild Cognitive Impairment and Alzheimer’s disease. Journal of the International Neuropsychological Society, 25(7), 688–698. https://doi.org/10.1017/S1355617719000316

39. Goldberg, T. E., Huey, E. D., & Devanand, D. P. (2020). Association of APOE e2 genotype with Alzheimer’s and non-Alzheimer’s neurodegenerative pathologies. Nature Communications, 11, 4727. https://doi.org/10.1038/s41467-020-18198-x

40. Gozdas, E., Fingerhut, H., Chromik, L. C., O’Hara, R., Reiss, A. L., & Hosseini, S. M. H. (2020). Focal white matter disruptions along the cingulum tract explain cognitive decline in amnestic mild cognitive impairment (aMCI). Scientific Reports, 10, 10213. https://doi.org/10.1038/s41598-020-66796-y

41. Harrison, J. R., Bhatia, S., Tan, Z. X., Mirza-Davies, A., Benkert, H., Tax, C. M. W., & Jones, D. K. (2020). Imaging Alzheimer’s genetic risk using diffusion MRI: A systematic review. NeuroImage: Clinical, 27, 102359. https://doi.org/10.1016/j.nicl.2020.102359

42. Heilbronner, S. R., & Haber, S. N. (2014). Frontal cortical and subcortical projections provide a basis for segmenting the cingulum bundle: Implications for neuroimaging and psychiatric disorders. Journal of Neuroscience, 34(30), 10041–10054. https://doi.org/10.1523/JNEUROSCI.5459-13.2014

43. Henson, R. N., Suri, S., Knights, E., Rowe, J. B., Kievit, R. A., Lyall, D. M., Chan, D., Eising, E., & Fisher, S. E. (2020). Effect of apolipoprotein E polymorphism on cognition and brain in the Cambridge Centre for Ageing and Neuroscience cohort. Brain and Neuroscience Advances, 4, 1–12. https://doi.org/10.1177/2398212820961704

44. Herbet, G., Zemmoura, I., & Duffau, H. (2018). Functional anatomy of the inferior longitudinal fasciculus: From historical reports to current hypotheses. Frontiers in Neuroanatomy, 12, 77. https://doi.org/10.3389/fnana.2018.00077

45. Herrup, K. (2015). The case for rejecting the amyloid cascade hypothesis. Nature Neuroscience, 18(6), 794–799. https://doi.org/10.1038/nn.4017

46. Hodgetts, C. J., Shine, J. P., Williams, H., Postans, M., Sims, R., Williams, J., Lawrence, A. D., & Graham, K. S. (2019). Increased posterior default mode network activity and structural connectivity in young adult APOE-ε4 carriers: A multimodal imaging investigation. Neurobiology of Aging, 73, 82–91. https://doi.org/10.1016/j.neurobiolaging.2018.08.026

47. Irfanoglu, M. O., Walker, L., Sarlls, J., Marenco, S., & Pierpaoli, C. (2012). Effects of image distortions originating from susceptibility variations and concomitant fields on diffusion MRI tractography results. NeuroImage, 61(1), 275–288. https://doi.org/10.1016/j.neuroimage.2012.02.054

48. Jacobs, H. I. L., Hedden, T., Schultz, A. P., Sepulcre, J., Perea, R. D., Amariglio, R. E., Papp, K. V., Rentz, D. M., Sperling, R. A., & Johnson, K. A. (2018). Structural tract alterations predict downstream tau accumulation in amyloid-positive older individuals. Nature Neuroscience, 21(3), 424–431. https://doi.org/10.1038/s41593-018-0070-z

49. Jagust, W. (2018). Imaging the evolution and pathophysiology of Alzheimer disease. Nature Reviews Neuroscience, 19(11), 687–700. https://doi.org/10.1038/s41583-018-0067-3

50. Jagust, W. J., & Mormino, E. C. (2011). Lifespan brain activity, β-amyloid, and Alzheimer’s disease. Trends in Cognitive Sciences, 15(11), 520–526. https://doi.org/10.1016/j.tics.2011.09.004

51. Jansen, W. J., Ossenkoppele, R., Knol, D. L., Tijms, B. M., Scheltens, P., Verhey, F. R. J., Visser, P. J., Aalten, P., Aarsland, D., Alcolea, D., Alexander, M., Almdahl, I. S., Arnold, S. E., Baldeiras, I., Barthel, H., van Berckel, B. N. M., Bibeau, K., Blennow, K., Brooks, D. J., … Zetterberg, H. (2015). Prevalence of cerebral amyloid pathology in persons without dementia: A meta-analysis. JAMA, 313(19), 1924–1938. https://doi.org/10.1001/jama.2015.4668

52. Jeurissen, B., Leemans, A., Jones, D. K., Tournier, J.-D., & Sijbers, J. (2011). Probabilistic fiber tracking using the residual bootstrap with constrained spherical deconvolution. Human Brain Mapping, 32(3), 461–479. https://doi.org/10.1002/hbm.21032

53. Jiang, L., Shing, N., Robin, J., Ladyka-Wojcik, N., Choi, A., Ryan, J. D., Barense, M. D., & Olsen, R. K. (2021). The association between visual discrimination and cognitive decline prior to clinical diagnosis. Alzheimer’s & Dementia, 17(S6), e057335. https://doi.org/10.1002/alz.057335

54. Jitsuishi, T., & Yamaguchi, A. (2021). Posterior precuneus is highly connected to medial temporal lobe revealed by tractography and white matter dissection. Neuroscience, 466, 173–185. https://doi.org/10.1016/j.neuroscience.2021.05.009

55. Jones, D. K., Christiansen, K. F., Chapman, R. J., & Aggleton, J. P. (2013). Distinct subdivisions of the cingulum bundle revealed by diffusion MRI fibre tracking: Implications for neuropsychological investigations. Neuropsychologia, 51(1), 67–78. https://doi.org/10.1016/J.NEUROPSYCHOLOGIA.2012.11.018

56. Jones, D. K., Horsfield, M. A., & Simmons, A. (1999). Optimal strategies for measuring diffusion in anisotropic systems by magnetic resonance imaging. Magnetic Resonance in Medicine, 42(3), 515–525. https://doi-org.abc.cardiff.ac.uk/10.1002/(SICI)1522-2594(199909)42:3<515::AID-MRM14>3.0.CO;2-Q

57. Jones, D. K., Knösche, T. R., & Turner, R. (2013). White matter integrity, fiber count, and other fallacies: The do’s and don’ts of diffusion MRI. NeuroImage, 73, 239–254. https://doi.org/10.1016/j.neuroimage.2012.06.081

58. Kantarci, K., Murray, M. E., Schwarz, C. G., Reid, R. I., Przybelski, S. A., Lesnick, T., Zuk, S. M., Raman, M. R., Senjem, M. L., Gunter, J. L., Boeve, B. F., Knopman, D. S., Parisi, J. E., Petersen, R. C., Jack, C. R., & Dickson, D. W. (2017). White-matter integrity on DTI and the pathologic staging of Alzheimer’s disease. Neurobiology of Aging, 56, 172–179. https://doi.org/10.1016/j.neurobiolaging.2017.04.024

59. Karch, C. M., Cruchaga, C., & Goate, A. M. (2014). Alzheimer’s disease genetics: From the bench to the clinic. Neuron, 83(1), 11–26. https://doi.org/10.1016/j.neuron.2014.05.041

60. Koelewijn, L., Lancaster, T. M., Linden, D., Dima, D. C., Routley, B. C., Magazzini, L., Barawi, K., Brindley, L., Adams, R., Tansey, K. E., Bompas, A., Tales, A., Bayer, A., & Singh, K. (2019). Oscillatory hyperactivity and hyperconnectivity in young APOE-ɛ4 carriers and hypoconnectivity in Alzheimer’s disease. ELife, 8, e36011. https://doi.org/10.7554/eLife.36011

61. Kor, D. Z. L., Jbabdi, S., Huszar, I. N., Mollink, J., Tendler, B. C., Foxley, S., Wang, C., Scott, C., Smart, A., Ansorge, O., Pallebage-Gamarallage, M., Miller, K. L., & Howard, A. F. D. (2022). An automated pipeline for extracting quantitative histological metrics for voxelwise MRI-histology comparisons. bioRxiv, 1-40. https://doi.org/10.1101/2022.02.10.479718

62. Lebel, C., Gee, M., Camicioli, R., Wieler, M., Martin, W., & Beaulieu, C. (2012). Diffusion tensor imaging of white matter tract evolution over the lifespan. NeuroImage, 60(1), 340–352. https://doi.org/10.1016/j.neuroimage.2011.11.094

63. Lee, A. C. H., Buckley, M. J., Gaffan, David., Emery, Tina., Hodges, J. R., & Graham, K. S. (2006). Differentiating the roles of the hippocampus and perirhinal cortex in processes beyond long-term declarative memory: A double dissociation in dementia. Journal of Neuroscience, 26(19), 5198–5203. https://doi.org/10.1523/JNEUROSCI.3157-05.2006

64. Lee, M. D., & Wagenmakers, E.-J. (2013). Bayesian cognitive modeling: A practical course (pp. xiii, 264). Cambridge University Press. https://doi.org/10.1017/CBO9781139087759

65. Leemans, A., Jeurissen, B., Sijbers, J., & Jones, D. K. (2009). ExploreDTI: A graphical toolbox for processing, analyzing, and visualizing diffusion MR data. Proceedings of the 17th Scientific Meeting, International Society for Magnetic Resonance in Medicine, 17, 3537.

66. Leemans, A., & Jones, D. K. (2009). The B-matrix must be rotated when correcting for subject motion in DTI data. Magnetic Resonance in Medicine, 61(6), 1336–1349. https://doi.org/10.1002/mrm.21890

67. Liu, C.-C., Kanekiyo, T., Xu, H., & Bu, G. (2013). Apolipoprotein E and Alzheimer disease: Risk, mechanisms, and therapy. Nature Reviews Neurology, 9(2), 106–118. https://doi.org/10.1038/nrneurol.2012.263

68. Lubben, N., Ensink, E., Coetzee, G. A., & Labrie, V. (2021). The enigma and implications of brain hemispheric asymmetry in neurodegenerative diseases. Brain Communications, 3(3), fcab211. https://doi.org/10.1093/braincomms/fcab211

69. Lupton, M. K., Medland, S. E., Gordon, S. D., Goncalves, T., MacGregor, S., Mackey, D. A., Young, T. L., Duffy, D. L., Visscher, P. M., Wray, N. R., Nyholt, D. R., Bain, L., Ferreira, M. A., Henders, A. K., Wallace, L., Montgomery, G. W., Wright, M. J., & Martin, N. G. (2018). Accuracy of inferred APOE genotypes for a range of genotyping arrays and imputation reference panels. Journal of Alzheimer’s Disease, 64(1), 49–54. https://doi.org/10.3233/JAD-171104

70. Ma, C., Wang, J., Zhang, J., Chen, K., Li, X., Shu, N., Chen, Y., Liu, Z., & Zhang, Z. (2017). Disrupted brain structural connectivity: Pathological interactions between genetic APOE ε4 status and developed MCI condition. Molecular Neurobiology, 54(9), 6999– 7007. https://doi.org/10.1007/s12035-016-0224-5

71. Maass, A., Berron, D., Harrison, T. M., Adams, J. N., La Joie, R., Baker, S., Mellinger, T., Bell, R. K., Swinnerton, K., Inglis, B., Rabinovici, G. D., Düzel, E., & Jagust, W. J. (2019). Alzheimer’s pathology targets distinct memory networks in the ageing brain. Brain, 142(8), 2492–2509. https://doi.org/10.1093/brain/awz154

72. Maechler, M., Rousseeuw, P., Croux, C., Todorov, V., Ruckstuhl, A., Salibian-Barrera, M., Verbeke, T., Koller, M., Conceicao, E. L., & Anna di Palma, M. (2021). robustbase: Basic robust statistics (Version 0.93-7) [Computer software]. http://CRAN.R-project.org/package=robustbase

73. Mason, E. J., Hussey, E. P., Molitor, R. J., Ko, P. C., Donahue, M. J., & Ally, B. A. (2017). Family history of Alzheimer’s disease is associated with impaired perceptual discrimination of novel objects. Journal of Alzheimer’s Disease, 57(3), 735–745. https://doi.org/10.3233/JAD-160772

74. MathWorks, Inc. (2015). MATLAB (Version R2015a) [Computer software]. https://uk.mathworks.com/

75. Mattsson, N., Palmqvist, S., Stomrud, E., Vogel, J., & Hansson, O. (2019). Staging β-amyloid pathology with amyloid positron emission tomography. JAMA Neurology, 76(11), 1319–1329. https://doi.org/10.1001/jamaneurol.2019.2214

76. Mayo, C. D., Mazerolle, E. L., Ritchie, L., Fisk, J. D., & Gawryluk, J. R. (2017). Longitudinal changes in microstructural white matter metrics in Alzheimer’s disease. NeuroImage: Clinical, 13, 330–338. https://doi.org/10.1016/j.nicl.2016.12.012

77. Mentink, L. J., Guimarães, J. P. O. F. T., Faber, M., Sprooten, E., Rikkert, M. G. M. O., Haak, K. V., & Beckmann, C. F. (2021). Functional co-activation of the default mode network in APOE ε4-carriers: A replication study. NeuroImage, 118304. https://doi.org/10.1016/j.neuroimage.2021.118304

78. Metzler-Baddeley, C., Jones, D. K., Steventon, J., Westacott, L., Aggleton, J. P., & O’Sullivan, M. J. (2012). Cingulum microstructure predicts cognitive control in older age and mild cognitive impairment. Journal of Neuroscience, 32(49), 17612–17619. https://doi.org/10.1523/JNEUROSCI.3299-12.2012

79. Metzler-Baddeley, C., Mole, J. P., Sims, R., Fasano, F., Evans, J., Jones, D. K., Aggleton, J. P., & Baddeley, R. J. (2019). Fornix white matter glia damage causes hippocampal gray matter damage during age-dependent limbic decline. Scientific Reports, 9, 1060. https://doi.org/10.1038/s41598-018-37658-5

80. Mishra, S., Blazey, T. M., Holtzman, D. M., Cruchaga, C., Su, Y., Morris, J. C., Benzinger, T. L. S., & Gordon, B. A. (2018). Longitudinal brain imaging in preclinical Alzheimer disease: Impact of APOE ε4 genotype. Brain, 141(6), 1828–1839. https://doi.org/10.1093/brain/awy103

81. Morey, R. D., & Rouder, J. N. (2018). BayesFactor: Computation of Bayes factors for common designs (Version 0.9.12-4.2) [Computer software]. https://CRAN.r-project.org/package=BayesFactor

82. Murray, E. A., Wise, S. P., & Graham, K. S. (2017). The evolution of memory systems: Ancestors, anatomy, and adaptations. Oxford University Press.

83. Oldmeadow, C., Holliday, E. G., McEvoy, M., Scott, R., Kwok, J. B. J., Mather, K., Sachdev, P., Schofield, P., & Attia, J. (2014). Concordance between direct and imputed APOE genotypes using 1000 Genomes data. Journal of Alzheimer’s Disease, 42(2), 391– 393. https://doi.org/10.3233/JAD-140846

84. Olsen, R. K., Yeung, L.-K., Noly-Gandon, A., D’Angelo, M. C., Kacollja, A., Smith, V. M., Ryan, J. D., & Barense, M. D. (2017). Human anterolateral entorhinal cortex volumes are associated with cognitive decline in aging prior to clinical diagnosis. Neurobiology of Aging, 57, 195–205. https://doi.org/10.1016/j.neurobiolaging.2017.04.025

85. Palmqvist, S., Schöll, M., Strandberg, O., Mattsson, N., Stomrud, E., Zetterberg, H., Blennow, K., Landau, S., Jagust, W., & Hansson, O. (2017). Earliest accumulation of β-amyloid occurs within the default-mode network and concurrently affects brain connectivity. Nature Communications, 8, 1214. https://doi.org/10.1038/s41467-017-01150-x

86. Panesar, S. S., Yeh, F.-C., Jacquesson, T., Hula, W., & Fernandez-Miranda, J. C. (2018). A quantitative tractography study into the connectivity, segmentation and laterality of the human inferior longitudinal fasciculus. Frontiers in Neuroanatomy, 12, 47. https://doi.org/10.3389/fnana.2018.00047

87. Parker, G. D. (2014). Robust processing of diffusion weighted image data [PhD, Cardiff University]. https://orca.cardiff.ac.uk/61622/

88. Parker, G. D., Marshall, D., Rosin, P. L., Drage, N., Richmond, S., & Jones, D. K. (2013). A pitfall in the reconstruction of fibre ODFs using spherical deconvolution of diffusion MRI data. NeuroImage, 65, 433–448. https://doi.org/10.1016/j.neuroimage.2012.10.022

89. Parker, G. D., Rosin, P. L., & Marshall, D. (2012). Automated segmentation of diffusion weighted MRI tractography. AVA / BMVA Meeting on Biological and Computer Vision, Spring (AGM) Meeting, Cambridge, United Kingdom.

90. Parvizi, J., Van Hoesen, G. W., Buckwalter, J., & Damasio, A. (2006). Neural connections of the posteromedial cortex in the macaque. Proceedings of the National Academy of Sciences, 103(5), 1563–1568. https://doi.org/10.1073/pnas.0507729103

91. Pasternak, O., Sochen, N., Gur, Y., Intrator, N., & Assaf, Y. (2009). Free water elimination and mapping from diffusion MRI. Magnetic Resonance in Medicine, 62(3), 717–730. https://doi.org/10.1002/mrm.22055

92. Pichet Binette, A., Theaud, G., Rheault, F., Roy, M., Collins, D. L., Levin, J., Mori, H., Lee, J. H., Farlow, M. R., Schofield, P., Chhatwal, J. P., Masters, C. L., Benzinger, T., Morris, J., Bateman, R., Breitner, J. C., Poirier, J., Gonneaud, J., Descoteaux, M., … PREVENT-AD Research Group. (2021). Bundle-specific associations between white matter microstructure and Aβ and tau pathology in preclinical Alzheimer’s disease. ELife, 10, e62929. https://doi.org/10.7554/eLife.62929

93. Postans, M., Hodgetts, C. J., Mundy, M. E., Jones, D. K., Lawrence, A. D., & Graham, K. S. (2014). Interindividual variation in fornix microstructure and macrostructure Is related to visual discrimination accuracy for scenes but not faces. Journal of Neuroscience, 34(36), 12121–12126. https://doi.org/10.1523/JNEUROSCI.0026-14.2014

94. Powell, J. L., Parkes, L., Kemp, G. J., Sluming, V., Barrick, T. R., & García-Fiñana, M. (2012). The effect of sex and handedness on white matter anisotropy: A diffusion tensor magnetic resonance imaging study. Neuroscience, 207, 227–242. https://doi.org/10.1016/j.neuroscience.2012.01.016

95. R Core Team. (2019). R: A language and environment for statistical computing (Version 3.6.0) [Computer software]. https://www.R-project.org/

96. Radmanesh, F., Devan, W. J., Anderson, C. D., Rosand, J., Falcone, G. J., & for the Alzheimer’s Disease Neuroimaging Initiative. (2014). Accuracy of imputation to infer unobserved APOE epsilon alleles in genome-wide genotyping data. European Journal of Human Genetics, 22(10), 1239–1242. https://doi.org/10.1038/ejhg.2013.308

97. Rajah, M. N., Wallace, L. M. K., Ankudowich, E., Yu, E. H., Swierkot, A., Patel, R., Chakravarty, M. M., Naumova, D., Pruessner, J., Joober, R., Gauthier, S., & Pasvanis, S. (2017). Family history and APOE4 risk for Alzheimer’s disease impact the neural correlates of episodic memory by early midlife. NeuroImage: Clinical, 14, 760–774. https://doi.org/10.1016/j.nicl.2017.03.016

98. Ranganath, C., & Ritchey, M. (2012). Two cortical systems for memory-guided behaviour. Nature Reviews Neuroscience, 13(10), 713–726. https://doi.org/10.1038/nrn3338

99. Reagh, Z. M., Ho, H. D., Leal, S. L., Noche, J. A., Chun, A., Murray, E. A., & Yassa, M. A. (2016). Greater loss of object than spatial mnemonic discrimination in aged adults. Hippocampus, 26(4), 417–422. https://doi.org/10.1002/hipo.22562

100. Reiman, E. M., Arboleda-Velasquez, J. F., Quiroz, Y. T., Huentelman, M. J., Beach, T. G., Caselli, R. J., Chen, Y., Su, Y., Myers, A. J., Hardy, J., Paul Vonsattel, J., Younkin, S. G., Bennett, D. A., De Jager, P. L., Larson, E. B., Crane, P. K., Keene, C. D., Kamboh, M. I., Kofler, J. K., … Jun, G. R. (2020). Exceptionally low likelihood of Alzheimer’s dementia in APOE2 homozygotes from a 5,000-person neuropathological study. Nature Communications, 11, 667. https://doi.org/10.1038/s41467-019-14279-8

101. Rieckmann, A., Van Dijk, K. R., Sperling, R. A., Johnson, K. A., Buckner, R. L., & Hedden, T. (2016). Accelerated decline in white matter integrity in clinically normal individuals at risk for Alzheimer’s disease. Neurobiology of Aging, 42, 177–188. https://doi.org/10.1016/j.neurobiolaging.2016.03.016

102. Riedel, B. C., Thompson, P. M., & Brinton, R. D. (2016). Age, APOE and sex: Triad of risk of Alzheimer’s disease. Journal of Steroid Biochemistry and Molecular Biology, 160, 134– 147. https://doi.org/10.1016/j.jsbmb.2016.03.012

103. Roe, J. M., Vidal-Piñeiro, D., Sørensen, Ø., Brandmaier, A. M., Düzel, S., Gonzalez, H. A., Kievit, R. A., Knights, E., Kühn, S., Lindenberger, U., Mowinckel, A. M., Nyberg, L., Park, D. C., Pudas, S., Rundle, M. M., Walhovd, K. B., Fjell, A. M., & Westerhausen, R. (2021). Asymmetric thinning of the cerebral cortex across the adult lifespan is accelerated in Alzheimer’s disease. Nature Communications, 12(1), 721. https://doi.org/10.1038/s41467-021-21057-y

104. RStudio Team. (2020). RStudio: Integrated development environment for R (Version 1.3.1093) [Computer software]. http://www.rstudio.com/

105. Scheltens, P., De Strooper, B., Kivipelto, M., Holstege, H., Chételat, G., Teunissen, C. E., Cummings, J., & van der Flier, W. M. (2021). Alzheimer’s disease. The Lancet, 397(10284), 1577–1590. https://doi.org/10.1016/S0140-6736(20)32205-4

106. Selkoe, D. J., & Hardy, J. (2016). The amyloid hypothesis of Alzheimer’s disease at 25 years. EMBO Molecular Medicine, 8(6), 595–608. https://doi.org/10.15252/emmm.201606210

107. Shine, J. P., Hodgetts, C. J., Postans, M., Lawrence, A. D., & Graham, K. S. (2015). APOE-ε4 selectively modulates posteromedial cortex activity during scene perception and short-term memory in young healthy adults. Scientific Reports, 5, 16322. https://doi.org/10.1038/srep16322

108. Silva, M. V. F., Loures, C., de M. G., Alves, L. C. V., de Souza, L. C., Borges, K. B. G., & Carvalho, M., das G. (2019). Alzheimer’s disease: Risk factors and potentially protective measures. Journal of Biomedical Science, 26(1), 33. https://doi.org/10.1186/s12929-019-0524-y

109. Smith, S. M., Jenkinson, M., Johansen-Berg, H., Rueckert, D., Nichols, T. E., Mackay, C. E., Watkins, K. E., Ciccarelli, O., Cader, M. Z., Matthews, P. M., & Behrens, T. E. J. (2006). Tract-based spatial statistics: Voxelwise analysis of multi-subject diffusion data. NeuroImage, 31(4), 1487–1505. https://doi.org/10.1016/j.neuroimage.2006.02.024

110. Smith, S. M., & Nichols, T. E. (2009). Threshold-free cluster enhancement: Addressing problems of smoothing, threshold dependence and localisation in cluster inference. NeuroImage, 44(1), 83–98. https://doi.org/10.1016/j.neuroimage.2008.03.061

111. Song, Z., Farrell, M. E., Chen, X., & Park, D. C. (2018). Longitudinal accrual of neocortical amyloid burden is associated with microstructural changes of the fornix in cognitively normal adults. Neurobiology of Aging, 68, 114–122. https://doi.org/10.1016/j.neurobiolaging.2018.02.021

112. Suri, S., Heise, V., Trachtenberg, A. J., & Mackay, C. E. (2013). The forgotten APOE allele: A review of the evidence and suggested mechanisms for the protective effect of APOE e2. Neuroscience & Biobehavioral Reviews, 37(10), 2878–2886. https://doi.org/10.1016/j.neubiorev.2013.10.010

113. Therriault, J., Benedet, A. L., Pascoal, T. A., Mathotaarachchi, S., Chamoun, M., Savard, M., Thomas, E., Kang, M. S., Lussier, F., Tissot, C., Parsons, M., Qureshi, M. N. I., Vitali, P., Massarweh, G., Soucy, J.-P., Rej, S., Saha-Chaudhuri, P., Gauthier, S., & Rosa-Neto, P. (2020). Association of apolipoprotein E ε4 with medial temporal tau independent of amyloid-β. JAMA Neurology, 77(4), 470–479. https://doi.org/10.1001/jamaneurol.2019.4421

114. Thiebaut de Schotten, M., ffytche, D. H., Bizzi, A., Dell’Acqua, F., Allin, M., Walshe, M., Murray, R., Williams, S. C., Murphy, D. G. M., & Catani, M. (2011). Atlasing location, asymmetry and inter-subject variability of white matter tracts in the human brain with MR diffusion tractography. NeuroImage, 54(1), 49–59. https://doi.org/10.1016/j.neuroimage.2010.07.055

115. Trejo-Lopez, J. A., Yachnis, A. T., & Prokop, S. (2021). Neuropathology of Alzheimer’s disease. Neurotherapeutics. https://doi.org/10.1007/s13311-021-01146-y

116. Tuch, D. S., Reese, T. G., Wiegell, M. R., Makris, N., Belliveau, J. W., & Wedeen, V. J. (2002). High angular resolution diffusion imaging reveals intravoxel white matter fiber heterogeneity. Magnetic Resonance in Medicine, 48(4), 577–582. https://doi.org/10.1002/mrm.10268

117. Tyrer, A., Gilbert, J. R., Adams, S., Stiles, A. B., Bankole, A. O., Gilchrist, I. D., & Moran, R. J. (2020). Lateralized memory circuit dropout in Alzheimer’s disease patients. Brain Communications, 2(2), fcaa212. https://doi.org/10.1093/braincomms/fcaa212

118. Verhagen, J., & Wagenmakers, E.-J. (2014). Bayesian tests to quantify the result of a replication attempt. Journal of Experimental Psychology: General, 143(4), 1457–1475. https://doi.org/10.1037/a0036731

119. Villeneuve, S., Rabinovici, G. D., Cohn-Sheehy, B. I., Madison, C., Ayakta, N., Ghosh, P. M., La Joie, R., Arthur-Bentil, S. K., Vogel, J. W., Marks, S. M., Lehmann, M., Rosen, H. J., Reed, B., Olichney, J., Boxer, A. L., Miller, B. L., Borys, E., Jin, L.-W., Huang, E. J., … Jagust, W. (2015). Existing Pittsburgh compound-B positron emission tomography thresholds are too high: Statistical and pathological evaluation. Brain, 138(7), 2020– 2033. https://doi.org/10.1093/brain/awv112

120. Vipin, A., Ng, K. K., Ji, F., Shim, H. Y., Lim, J. K. W., Pasternak, O., Zhou, J. H., & for the Alzheimer’s Disease Neuroimaging Initiative. (2019). Amyloid burden accelerates white matter degradation in cognitively normal elderly individuals. Human Brain Mapping, 40(7), 2065–2075. https://doi.org/10.1002/hbm.24507

121. Wakana, S., Caprihan, A., Panzenboeck, M. M., Fallon, J. H., Perry, M., Gollub, R. L., Hua, K., Zhang, J., Jiang, H., Dubey, P., Blitz, A., van Zijl, P., & Mori, S. (2007). Reproducibility of quantitative tractography methods applied to cerebral white matter. NeuroImage, 36(3), 630–644. https://doi.org/10.1016/j.neuroimage.2007.02.049

122. Weise, C. M., Chen, K., Chen, Y., Kuang, X., Savage, C. R., & Reiman, E. M. (2018). Left lateralized cerebral glucose metabolism declines in amyloid-β positive persons with mild cognitive impairment. NeuroImage: Clinical, 20, 286–296. https://doi.org/10.1016/j.nicl.2018.07.016

123. Winkler, A. M., Ridgway, G. R., Webster, M. A., Smith, S. M., & Nichols, T. E. (2014). Permutation inference for the general linear model. NeuroImage, 92, 381–397. https://doi.org/10.1016/j.neuroimage.2014.01.060

124. Wolf, D., Fischer, F. U., Scheurich, A., Fellgiebel, A., & for the Alzheimer’s Disease Neuroimaging Alzheimer’s Disease Neuroimaging Initiative. (2015). Non-linear association between cerebral amyloid deposition and white matter microstructure in cognitively healthy older adults. Journal of Alzheimer’s Disease, 47(1), 117–127. https://doi.org/10.3233/JAD-150049

125. Yeh, C.-H., Jones, D. K., Liang, X., Descoteaux, M., & Connelly, A. (2021). Mapping structural connectivity using diffusion MRI: Challenges and opportunities. Journal of Magnetic Resonance Imaging, 53(6), 1666–1682. https://doi.org/10.1002/jmri.27188

126. Yu, J., Lam, C. L. M., & Lee, T. M. C. (2017). White matter microstructural abnormalities in amnestic mild cognitive impairment: A meta-analysis of whole-brain and ROI-based studies. Neuroscience & Biobehavioral Reviews, 83, 405–416. https://doi.org/10.1016/j.neubiorev.2017.10.026

127. Zhao, J., Thiebaut de Schotten, M., Altarelli, I., Dubois, J., & Ramus, F. (2016). Altered hemispheric lateralization of white matter pathways in developmental dyslexia: Evidence from spherical deconvolution tractography. Cortex, 76, 51–62. https://doi.org/10.1016/j.cortex.2015.12.004

128. Ziontz, J., Adams, J. N., Harrison, T. M., Baker, S. L., & Jagust, W. J. (2021). Hippocampal connectivity with retrosplenial cortex is linked to neocortical tau accumulation and memory function. Journal of Neuroscience, 41(42), 8839–8847. https://doi.org/10.1523/JNEUROSCI.0990-21.2021

