## Supplementary Tables & Figures for "Tract-specific white matter microstructure alterations among young adult *APOE* ε4 carriers: A replication and extension study"

**Supplementary Table 1**

*Comparison of Basic Sample Characteristics Used in the Current Study and in Hodgetts et al. (2019)*

| | Current Study | | Hodgetts et al. (2019) | |
|---|---|---|---|---|
| | *APOE* ε4+ | *APOE* ε4- | *APOE* ε4+ | *APOE* ε4- |
| Total *n* | 40 | 88 | 15 | 15 |
| *APOE* genotype split (*n*) | 4 ε2/ε4, 33 ε3/ε4, 3 ε4/ε4 | 4 ε2/ε2, 14 ε2/ε3, 70 ε3/ε3 | 1 ε2/ε4, 14 ε3/ε2, 0 ε4/ε4 | 0 ε2/ε2, 5 ε2/ε3, 10 ε3/ε3 |
| Age (years; *M* ± *SD*) | 23.9 ±3.3 | 23.7 ±3.7 | 19.7 ±0.84 | 19.7 ±0.89 |
| Sex (Males/Females; *n*) | 12/28 | 30/58 | 1/14 | 1/14 |

*Note*. Abbreviations: *APOE* ε4+ = *APOE* ε4 carrier, *APOE* ε4- = *APOE* ε4 non-carrier, *M* = mean, *n* = number of participants, *SD* = standard deviation.

**Supplementary Table 2**

*Comparison of the MRI Scan Parameters Used in the Current Study and in Hodgetts et al. (2019)*

| | Current Study | Hodgetts et al. (2019) |
|---|---|---|
| *Diffusion-weighted sequence* | | |
| TE | 89ms | 87ms |
| Voxel dimensions | 2.4 x 2.4 x 2.4mm | 2.4 x 2.4 x 2.4mm |
| FOV | 230 x 230mm | 230 x 230mm |
| Acquisition matrix | 96 x 96 | 96 x 96 |
| Slices | 60 (aligned AC/PC) with 2.4mm thickness, no gap | 60 slices (oblique axial) with 2.4mm thickness, no gap |
| Diffusion gradients | 30 isotropic directions (b = 1200 s/mm$^2$), 3 non-diffusion images (b = 0 s/mm$^2$) | 30 isotropic directions (b = 1200 s/mm$^2$), 3 non-diffusion images (b = 0 s/mm$^2$) |
| *Structural sequence* | | |
| TR | 7.8s | 7.8s |
| TE | 3s | 3s |
| Voxel dimensions | 1 x 1 x 1mm | 1 x 1 x 1mm |
| FOV | 256 x 256 x 168mm - 256 x 256 x 180mm | 256 x 256 x 176mm |
| Acquisition matrix | 256 x 256 x 168 – 256 x 256 x 180 | 256 x 256 x 176 |
| Flip angle | 20° | 20° |

*Note*. The structural and diffusion-weighted sequences are remarkably similar across studies. In addition, the sequences were run on the same MRI system, thereby reducing the likelihood that differences in data acquisition can readily account for any discrepancies in results. Abbreviations: AC/PC = anterior commissure/posterior commissure, FOV = field of view, TE = echo time, TR = repetition time.

**Supplementary Figure 1**

*Flowchart of the Progress from Data Availability to Inclusion in the Study*

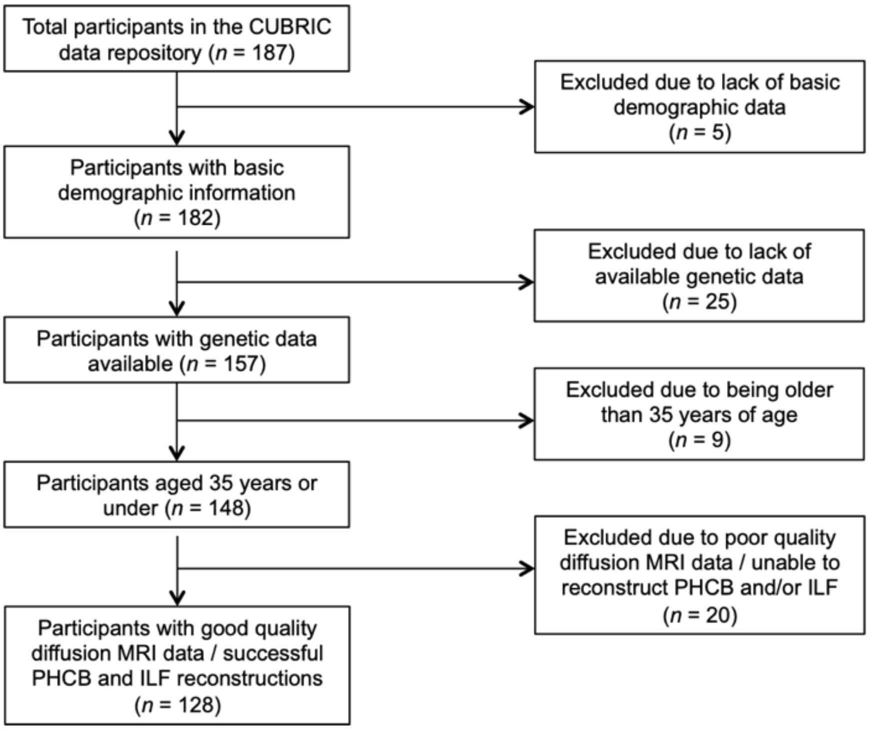


*Note.* Of the total number of participants in the CUBRIC data repository, approximately two-thirds (68.45%) were included in the final sample reported here. All exclusions are reported in the main body of the manuscript. Abbreviations: PHCB = parahippocampal cingulum bundle, ILF = inferior longitudinal fasciculus.
